# Computational Convergence of Adaptive Immunity and Artificial Intelligence

**DOI:** 10.64898/2026.02.03.703525

**Authors:** Sai T. Reddy

## Abstract

The adaptive immune system and modern artificial intelligence (AI) have independently converged on identical computational strategies for solving the recognition and generalization problem: learning to identify which inputs should be associated with which outputs, such that performance extends to novel instances never seen during training. We demonstrate that four major AI architectural innovations can be derived from immunological first principles. Two represent exact mathematical equivalences: the softmax function in transformer attention is the Boltzmann distribution governing antibody-antigen binding; the InfoNCE loss in contrastive learning is the negative log of clonal selection probability. Two represent strategic convergences: the pre-training/fine-tuning/RLHF training hierarchy parallels the germline/somatic hypermutation/T follicular helper stages of antibody affinity maturation; retrieval-augmented generation mirrors the long-lived plasma cell and memory B cell dual memory system. Critically, these derivations run from immunology to AI, not the reverse—the mathematical structures of transformer attention and contrastive learning could have been derived from immunological first principles before their empirical discovery in AI. The framework generates testable predictions for immune specificity foundation models and suggests that certain computational strategies may be fundamental to any system solving recognition and generalization under resource constraints. Modern AI has independently converged on computational strategies that the adaptive immune system discovered through 500 million years of evolution.

**One-sentence summary:** The mathematics of transformer attention and contrastive learning can be derived from the biophysics of immune recognition.

## Introduction

Artificial intelligence and the adaptive immune system solve the same fundamental problem: learning to determine which inputs should be associated with which outputs based on patterns of similarity, such that performance extends to novel instances not seen during training. In AI, this manifests as selective attention (which tokens matter for this prediction?), contrastive discrimination (which examples are similar versus dissimilar?), and retrieval (which stored items match this query?) - all requiring generalization beyond training data (*1–3*). In immunology, this is molecular recognition: does this antibody bind this antigen? Should this T cell respond to this peptide-MHC complex? The immune system must respond to novel pathogens it has never encountered, using receptors generated through stochastic recombination (*4, 5*). Both involve similarity-based matching across vast combinatorial spaces, learning from limited examples, and generalizing to novel instances under severe resource constraints.

This paper presents evidence that both systems have independently converged on the same computational strategies. The correspondences we identify are not vague analogies. For transformer attention (*1*) and contrastive learning (*2*), they are mathematical identities: the softmax function is the Boltzmann distribution - a relationship established in statistical mechanics over a century ago (*6*) and recognized in machine learning contexts (*7*). Recent work has connected transformer attention to modern Hopfield networks (*8*), demonstrating that attention mechanisms can be understood through the lens of energy-based associative memory. The mathematical equivalences are not in dispute.

What has not been established is why these mathematical structures appear in both domains, and whether they can be derived from one domain to the other. Our contribution is not to note that softmax equals Boltzmann, but to show that the complete transformer architecture - including the query-key-value decomposition, multi-head parallelism, and cross-attention asymmetry - emerges naturally from the biophysics of antibody-antigen binding (*9, 10*). Similarly, the InfoNCE loss is mathematically equivalent to the negative log of immune selection probability, and we demonstrate that the entire contrastive learning framework can be derived from clonal selection dynamics (*11, 12*). For training hierarchies and memory architectures, we identify strategic convergences: the same computational structure solving the same optimization problem, with analogous functional roles at each stage (*13–17*).

Critically, these derivations run from immunology to AI architecture, not the reverse. An immunologist reasoning from first principles about binding thermodynamics in 2012 - five years before “Attention Is All You Need” (*1*) - could have derived the attention mechanism from the biophysics of antibody-antigen binding. The equations governing thymic selection (*18, 19*) could have motivated the InfoNCE loss before contrastive learning emerged as a field. Prior work applies AI architectures to immune prediction; we argue that immunology provides a principled foundation from which modern AI architectures can be independently derived.

We term this phenomenon **computational convergence**: the independent discovery of equivalent computational strategies by biological evolution and human engineering, driven by shared constraints on solving recognition and generalization. Both systems face astronomically large input spaces: theoretical adaptive immune receptor sequence diversity is estimated at 10^13^ - 10^18^ (*20, 21*), while the potential antigen space exceeds 10^15^ distinct epitopes (*22, 23*), and text sequences for language are effectively infinite. Both learn from sparse data: the immune system from developmental exposure and potentially thousands of distinct pathogen encounters over a lifetime (*24*); AI from training corpora that, while massive, represent a vanishing fraction of possible inputs (*25*). Both must generalize to novel instances never seen during training (*26, 27*). Both face tradeoffs between maintaining existing knowledge and adapting to new patterns (*28, 29*). Both operate under resource constraints requiring compression and prioritization.

Shared constraints yield shared solutions - convergent evolution at the computational level. Convolutional neural networks trained on images independently learn edge detectors matching the Gabor filters of primary visual cortex (*30*). Bats evolved echolocation using the same pulse-echo ranging as engineered sonar (*31*). In both cases, biological and engineered systems converged on identical computations because they faced identical problems. The convergences we document follow this pattern: transformer attention and antibody binding both solve the problem of selective, similarity-weighted information aggregation under pressure to generalize.

We present four convergences between AI architectures and adaptive immunity (**Table 1**). The AI systems we examine - transformer attention, contrastive learning objectives, multi-stage training with reinforcement learning, and retrieval-augmented generation - emerged primarily in large language models but extend across vision, multimodal systems, and protein modeling (*32–35*). The mathematical correspondences apply across these domains.

**Table 1.** Computational Convergences Between AI and Adaptive Immunity.

| Convergence | AI Architecture (Year) | Immunological Principle (Year) | Mathematical Core | Type |
| --- | --- | --- | --- | --- |
| 1. Attention | Softmax attention (2017) | Boltzmann distribution (1877) + Ab-Ag structure (1986) | $\exp(s)/\sum \exp(s)$ | **Exact equivalence** |
| 2. Contrastive | InfoNCE loss (2018) | Clonal selection (1959) + AIRE (2002) | $-\log P(\text{select positive})$ | **Exact equivalence** |
| 3. Training | Pre-train/Fine-tune/R LHF (2017–2022) | Germline/SHM/Tfh (1959–2000) | Three-stage hierarchy | Strategic convergence |
| 4. Memory | RAG (2020) | Plasma cells + Memory B cells (1996) | Dual architecture | Strategic convergence |

## Results

### Overview

We distinguish two types of convergence. Exact mathematical equivalence (Convergences 1 and 2): the equations are identical - softmax is Boltzmann; InfoNCE is selection probability. Strategic convergence (Convergences 3 and 4): the computational structure is the same, with components serving analogous functional roles.

#### Convergence 1: Transformer Attention from Antibody-Antigen Binding

##### Immunological starting point

Antibody-antigen binding involves selective contact between amino acid residues (*9, 10, 36*). When an antibody paratope (binding site) engages an antigen epitope (target region), specific residue pairs come into contact, and the probability of each contact depends on its energetic favorability (*37*). At thermal equilibrium, the probability that paratope residue *i* contacts epitope residue *j* follows the Boltzmann distribution - a fundamental result from statistical mechanics (*6*):

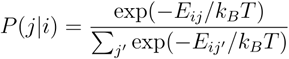

where *E*_*ij*_ is the interaction energy, *K*_*B*_ is Boltzmann’s constant, and *T* is temperature. Lower energy contacts (more favorable) have higher probability; higher energy contacts (less favorable) have lower probability. This is textbook biophysics - the only assumption is thermal equilibrium.

##### Mathematical equivalence

Define a dimensionless compatibility score as the negative energy scaled by thermal energy: *s*_*ij*_ *=*− *E*_*ij*_ /*k*_*B*_ *T* . The Boltzmann distribution becomes:

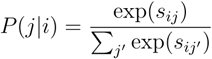

This is the softmax function - the same function that defines attention weights in transformers (*1, 7*). The equivalence is exact, not approximate. What immunology calls a “contact probability” and what AI calls an “attention weight” are the same mathematical object:

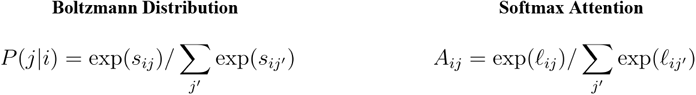

##### Query-key-value structure

Transformer attention computes compatibility as dot products between learned query and key vectors: *s*_*ij*_ *= q*_*i*_ · *k*_*j*_ (*1*). This has direct structural interpretation in molecular recognition. The paratope residue *i* can be represented by a feature vector (its “query”); the epitope residue *j* can be represented by a feature vector (its “key”). Their compatibility - how favorably they interact - is captured by the similarity of these vectors. The value vectors correspond to information transferred upon contact. The attention output aggregates contributions weighted by contact probability:

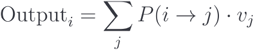

This is precisely how binding interfaces work: each paratope residue integrates information from epitope residues it contacts, weighted by contact strength (*10*).

##### Multi-head attention and CDR loops

Antibodies have six complementarity-determining region (CDR) loops - three on the heavy chain (CDRH1-3) and three on the light chain (CDRL1-3) (*38*). These loops form the paratope and make contact with the epitope. Crucially, different CDR loops often engage different regions of the epitope and contribute different interaction types (electrostatic, hydrophobic, hydrogen bonding) (*39*). This parallels multi-head attention: *H* parallel attention heads with independent parameters, each specializing for different patterns, with outputs concatenated and combined (*1*). The architectural principle is shared: parallel recognition modules with independent parameters that integrate their contributions.

##### Temperature and scaling

The Boltzmann distribution includes temperature *T*, which controls distribution sharpness. At low temperature, probability concentrates on lowest-energy contacts; at high temperature, the distribution becomes uniform. Transformer attention includes a scaling factor 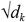 serving the same function, sometimes written explicitly as temperature τ (*40*). The correspondence is exact: Boltzmann *T* ↔attention τ; low *T* ↔sharp attention; high *T* ↔diffuse attention.

##### The counterfactual derivation

An immunologist in 2012 could have derived transformer attention: (1) start with binding physics - contact probabilities follow Boltzmann (*6*); (2) define compatibility scores as *s*_*ij*_ *=*− *E*_*ij*_ /*k*_*B*_ *T*; (3) recognize this is the softmax function (*7*); (4) parameterize compatibility as dot products of feature vectors (query-key); (5) add parallel channels for different interaction types (multi-head) (*38*); (6) aggregate information weighted by contact probability. The result is:

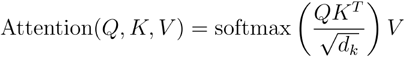

This is the transformer attention equation from Vaswani et al. (*1*). It could have been written down from immunological first principles five years before its publication. The derivation requires only undergraduate physics (Boltzmann, 1877) and structural immunology (CDR loops, Chothia & Lesk, 1987 (*38*); contact interfaces, Amit et al., 1986 (*10*)). No machine learning knowledge is needed.

Full mathematical details in **Supplementary Note S1**.

#### Convergence 2: Contrastive Learning from Clonal Selection

##### Immunological starting point

The adaptive immune system faces a fundamental discrimination problem: receptors must respond to foreign antigens (pathogens) while ignoring self-antigens (the body’s own molecules) (*11, 12*). This discrimination is enforced through selection - cells with receptors that bind foreign antigens are amplified; cells that bind self-antigens are eliminated (*18, 41*).

Consider T cell development in the thymus (*19, 42*). Each developing T cell (thymocyte) expresses a T cell receptor (TCR) generated by V(D)J recombination (*4, 5*). This TCR is tested against peptides presented on MHC molecules: **positive selection** ensures TCRs bind self-MHC with moderate affinity (MHC restriction) (*43*); **negative selection** eliminates TCRs that bind self-peptide-MHC too strongly (preventing autoimmunity) (*18, 44*). The result is a repertoire that recognizes foreign peptides on self-MHC while tolerating self-peptides. This is a contrastive discrimination task: distinguish the positive (foreign) from negatives (self) based on binding affinity.

##### Mathematical equivalence

Consider a receptor *r* encountering a set of candidate antigens: one foreign antigen *f* (the “positive”) and *M* self-antigens *s*_*1*_, …, *s*_*M*_ (the “negatives”). Let *a(r,p)* denote binding affinity between receptor *r* and antigen *p*. Under competitive selection, the probability that receptor *r* selects the foreign antigen follows a softmax over affinities (*45*):

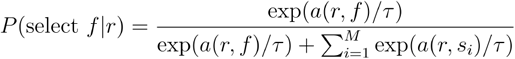

where *τ* controls selection stringency.

In contrastive learning, an anchor *x* is paired with a positive *x*^+^ and negatives 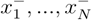 (*2, 46*). Let sim*(z,z*^′^) denote similarity between embeddings. The probability of correctly selecting the positive is:

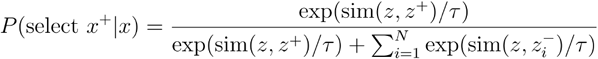

These equations are identical. What immunology calls “selection probability” and what AI calls “softmax over similarities” are the same mathematical object. The InfoNCE loss - the standard objective for contrastive learning (*2*) - is simply the negative log of this probability:

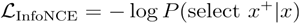

This is the negative log-likelihood of correct discrimination. Minimizing InfoNCE maximizes the probability of selecting the positive over negatives - exactly what thymic selection accomplishes for T cells.

##### Hard negatives and AIRE

A key insight from contrastive learning is that hard negatives - negatives similar to the positive - improve discrimination (*47, 48*). Easy negatives provide little training signal; the model needs challenging examples.

The immune system implements exactly this through AIRE (Autoimmune Regulator) (*49*). AIRE is a transcription factor expressed in thymic medullary epithelial cells that drives ectopic expression of tissue-restricted antigens - proteins normally found only in specific organs (pancreas, thyroid, etc.) (*19, 50*). Why would the thymus express pancreatic proteins? Because these are the hard negatives. A T cell that cross-reacts with insulin (pancreas-specific) would escape selection unless insulin is presented in the thymus. AIRE ensures developing T cells are tested against the most challenging self-antigens.

AIRE deficiency causes autoimmune polyendocrine syndrome (APS-1), with autoimmunity against multiple organs - the biological equivalent of training without hard negatives: the system fails to learn fine-grained discrimination.

##### Temperature and selection stringency

The temperature parameter τ controls sharpness of selection (*52*). Low τ (stringent selection) means small affinity differences produce large probability differences - only the highest-affinity interaction dominates. High τ (permissive selection) means probabilities approach uniform - multiple candidates have similar selection probability. In thymic selection, this corresponds to how much better must a receptor bind foreign versus self to survive (*53*). In contrastive learning, τ controls the same tradeoff (*40*). This is the same parameter controlling the same mathematical tradeoff in the same equation.

##### Germinal center selection

The same mathematics applies to B cell selection in germinal centers (*54, 55*). B cells compete for T follicular helper (Tfh) cell help based on their ability to capture and present antigen (*17*). Higher-affinity B cells capture more antigen, present more peptide-MHC, and receive more Tfh help. This is softmax competition over the batch of competing B cells—the same equation describing competitive selection in contrastive learning batches.

##### The counterfactual derivation

An immunologist in 2012 could have derived InfoNCE: (1) model selection as softmax over affinities (Boltzmann-like competition); (2) define the training objective as maximizing P(select foreign | receptor); (3) take the negative log; (4) add hard negatives based on AIRE biology (*49*); (5) tune temperature for selection stringency. The result is the InfoNCE loss (*2*), derivable from Burnet’s clonal selection theory (1959) (*11*) decades before its invention in AI.

Full mathematical details in **Supplementary Note S2**.

#### Convergence 3: Training Hierarchy from Affinity Maturation

Both systems face a multi-scale optimization problem: (1) What general capabilities should the system have? (2) How should it specialize for a specific challenge? (3) How should it calibrate the magnitude and style of response? A single optimization process struggles to handle all three scales efficiently. Both systems converged on the same solution: a three-stage hierarchy (**Table 2**).

**Table 2.** Three-Stage Training Hierarchy.

| Stage | Immune System | AI Training | Function | Mathematical Status |
| --- | --- | --- | --- | --- |
| 1 | Germline V(D)J genes | Pre-training | Optimized prior | Structural correspondence |
| 2 | Somatic hypermutation ($\mu \approx 10^{-3}$) | Fine-tuning ($\eta \approx 10^{-3}$) | Local adaptation | Structural correspondence |
| 3 | Tfh selection | RLHF | Preference calibration | Exact equivalence (softmax) |

**Table 3.**
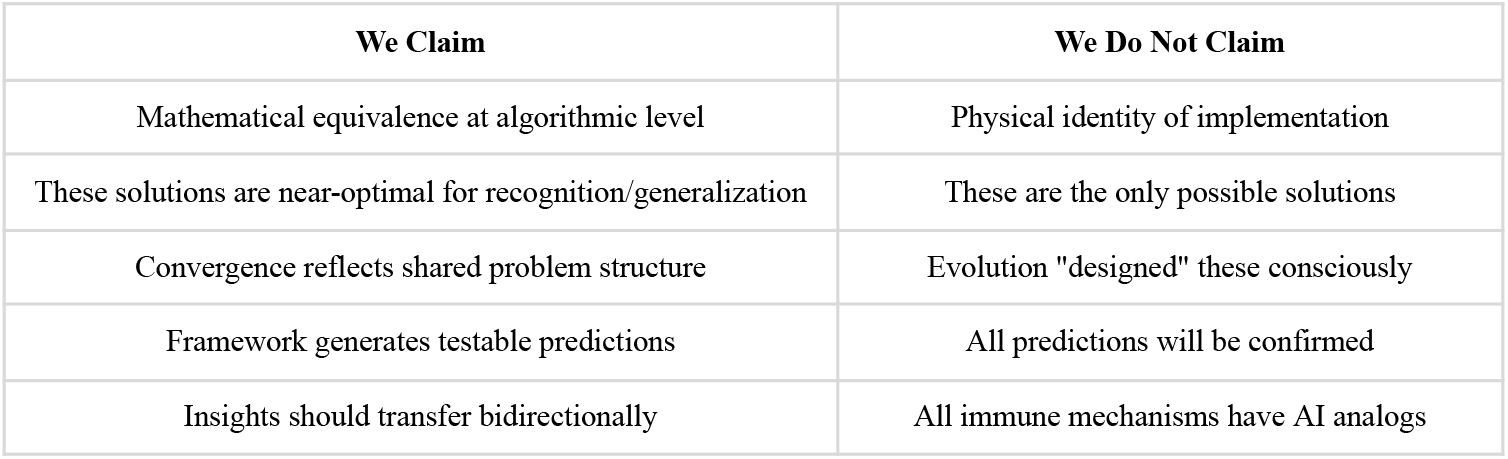
What We Claim vs. Do Not Claim.

| We Claim | We Do Not Claim |
| --- | --- |
| Mathematical equivalence at algorithmic level | Physical identity of implementation |
| These solutions are near-optimal for recognition/generalization | These are the only possible solutions |
| Convergence reflects shared problem structure | Evolution "designed" these consciously |
| Framework generates testable predictions | All predictions will be confirmed |
| Insights should transfer bidirectionally | All immune mechanisms have AI analogs |

##### Stage 1: Germline ↔Pre-training (Optimized Prior)

Antibodies are assembled from V (variable), D (diversity), and J (joining) gene segments (*4, 13*). These germline genes are not random - they have been optimized by millions of years of evolution (*59, 60*). The germline encodes a prior distribution over antibody space:

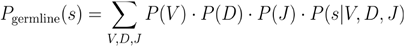

where gene usage frequencies reflect evolutionary optimization (*57*). This distribution is non-uniform, biased toward sequences that fold into stable antibody structures and bind common pathogen motifs.

A pre-trained language model similarly encodes patterns from training data (*58, 59*). The pre-trained weights define a conditional distribution 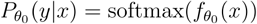 that reflects grammar, syntax, world knowledge, and reasoning patterns. Both germline and pre-training create optimized priors - starting points much better than random, shaped by extensive optimization on diverse data.

##### Stage 2: SHM ↔Fine-tuning (Local Adaptation)

When a B cell enters a germinal center, it undergoes somatic hypermutation (SHM) (*14*). The enzyme AID introduces point mutations at rate μ ≈ 10^−3^ per base pair per division - roughly one million times faster than normal replication error (*60*). SHM creates variants around the germline starting point; combined with selection, this implements local optimization.

Fine-tuning adapts pre-trained weights θ_0_ to a specific task (*15*): *θ*_*t*+1_ = *θ*_*t*_ − *η* ∇_*θ*_ ℒ (*θ*_*t*_), where η ≈ 10^−3^ to 10^−5^ is the learning rate - deliberately small to preserve pre-trained knowledge. Both make small local changes from good starting points. Large changes would destroy what makes the starting point valuable.

The magnitude similarity is striking: both use step sizes on the order of 10^−3^. This may reflect a shared constraint - the “Goldilocks zone” for adaptation from an optimized prior. Additionally, mutations in SHM concentrate in CDRs (binding regions) while sparing framework regions (*61*); common practice in fine-tuning uses different learning rates for different layers (*62*). Adaptation focuses where it matters most.

##### Stage 3: Tfh ↔RLHF (Preference Calibration)

After SHM generates variants, which survive? T follicular helper (Tfh) cells determine this (*17, 63*). B cells compete for Tfh help based on antigen presentation quality—a proxy for affinity, not ground truth (*64*). Critically, the B cell receives “you’re doing well,” not “your affinity is X.”

RLHF (Reinforcement Learning from Human Feedback) similarly aligns model behavior with human preferences (*16, 65*). Human raters compare responses (“A is better than B”), a reward model learns to predict preferences, and the language model optimizes toward higher reward. Critically, humans provide preference feedback—not the exact correct answer.

Both stages use preference feedback because ground truth is unavailable or impractical. Comparing “A vs. B” is easier than specifying the optimal answer. The selection mathematics is identical—softmax competition over candidates based on quality/reward scores:

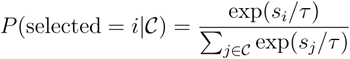

where *s*_*i*_ is quality proxy (Tfh) or reward model score (RLHF).

###### Why three stages?

Different objectives require different feedback types with different costs. Stage 1 uses cheap, abundant implicit feedback (survival/prediction). Stage 2 uses task-specific explicit feedback (affinity/task loss). Stage 3 uses expensive preference feedback (limited scale). If different objectives require different feedback types, and feedback types have different costs, staging becomes natural: use cheap feedback first, task-specific feedback second, expensive preference feedback last.

Why not fewer stages? **One stage** cannot efficiently combine general learning with task-specific optimization—you cannot do RLHF-style training with trillions of examples. **Two stages** lack calibration: a model can be highly capable but generate harmful outputs or have wrong tone (*66*). Stage 3 provides calibration that capability optimization alone cannot achieve.

Full mathematical details in **Supplementary Note S3**.

#### Convergence 4: Dual Memory from Immune Memory Architecture

##### The problem

Both systems face a fundamental memory problem: how to remember vast amounts of information while responding quickly to current challenges? The immune system must remember every pathogen encountered—potentially thousands over a lifetime (*24*)—while maintaining the ability to respond immediately to reinfection. Storing detailed responses in continuously active form would be metabolically prohibitive (*67*). An AI system must access vast knowledge while generating responses in real-time. Encoding all knowledge in parameters would require impossibly large models and cause catastrophic forgetting during updates (*28*).

Both systems converged on the same solution: **dual memory architecture**—two complementary memory systems with different properties, combined with similarity-based retrieval (*68, 69*).

##### Immune memory: Plasma cells and memory B cells

The adaptive immune system maintains two distinct populations of antigen-experienced B cells (*70, 71*):

###### Long-lived plasma cells (LLPCs)

Reside in bone marrow niches; constitutively secrete antibodies without requiring reactivation; provide immediate, always-on protection; limited number (∼10^4^–10^5^ niches); metabolically expensive to maintain (*67*).

###### Memory B cells

Circulate in blood and lymphoid tissues; quiescent until antigen re-encounter; require activation to produce antibody; can be maintained in large numbers (∼10^7^–10^8^ cells); metabolically inexpensive when dormant (*72, 73*).

This is a dual memory system: always-on protection from plasma cells, augmented by on-demand recall from memory B cells.

##### RAG: Parametric and retrieved memory

Retrieval-Augmented Generation (*3*) similarly maintains two memory systems:

###### Parametric memory (model weights)

Knowledge encoded in neural network parameters; always available during generation; provides immediate responses; limited by model capacity; expensive to update (requires retraining).

###### Retrieved memory (external knowledge base)

Documents stored in searchable database; retrieved on-demand based on query (*74*); requires retrieval step before use; can store virtually unlimited information; cheap to update (just add documents).

#### Similarity-based retrieval

Both systems retrieve memories based on similarity between query and stored items. Memory B cells compete for activation based on BCR affinity (*75*):

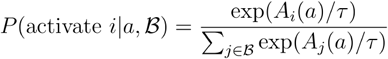

where *A*_*i*_ (*a*) is the affinity of cell *i*’s BCR for antigen *a*. Document retrieval follows the same form (*80*):

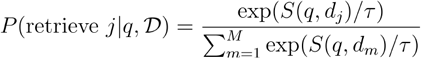

where *S*(*q,d*_*j*_ ) is embedding similarity between query and document. The mathematical structure is identical—the same Boltzmann-form softmax appearing in Convergences 1, 2, and 3.

#### Resource allocation

Both systems must decide how to allocate limited “always-on” resources versus larger “on-demand” stores (*76*). Bone marrow niches are limited—which specificities get permanent slots? Evidence suggests allocation depends on frequency (commonly encountered pathogens) and severity (dangerous pathogens) (*29, 77*). Similarly, RAG systems encode frequently used, time-critical knowledge in parameters while storing the long tail in retrieval.

The fundamental tradeoff: if everything were plasma cells/parametric, metabolic/compute cost would be prohibitive. If everything were memory B cells/retrieved, response latency would be too slow (*78*). The dual architecture resolves this by partitioning memory by access pattern: frequently needed, time-critical → always-on; rarely needed, can tolerate delay → on-demand.

Full mathematical details in **Supplementary Note S4**.

### Testable Predictions

The framework generates falsifiable predictions for immune specificity foundation models (ISFMs)—transformer architectures trained with contrastive objectives on antibody-antigen and TCR-pMHC data.

#### Predictions requiring discriminative capability

- Attention weights should correlate with experimental binding contacts
- Different attention heads should specialize for different interaction types
- Model-predicted affinity rankings should match experimental rankings

#### Predictions requiring generalization

- Hard negative mining should improve out-of-distribution generalization
- Germline-biased initialization should improve sample efficiency
- Learned temperature should converge to biologically interpretable values

#### Predictions requiring scaling characterization

- Loss should follow power-law scaling with training data
- Data diversity should matter more than quantity for generalization

### Falsification criteria

If a well-performing ISFM shows no correlation between attention and experimental contacts, this would challenge the claim that attention implements Boltzmann-weighted binding. If hard negative mining provides no out-of-distribution benefit, this would challenge the AIRE correspondence.

## Discussion

We have shown that four core AI architectures can be derived from immunological first principles. Two are exact mathematical equivalences: softmax attention is the Boltzmann distribution (*6*); InfoNCE is the negative log of selection probability (*2, 11*). Two are strategic convergences: the three-stage training hierarchy mirrors affinity maturation (*13–17*); dual memory architecture mirrors the plasma cell and memory B cell system (*67–73*).

These correspondences are not retrospective analogies. The mathematics of attention and contrastive learning is derivable from immunological first principles—from the biophysics of antibody-antigen binding (*9, 10*) and the dynamics of clonal selection (*11, 12*). An immunologist in 2012, armed with the Boltzmann distribution and *Janeway’s Immunobiology*, could have derived the attention mechanism five years before its appearance in AI (and published “Antibody-Antigen is All You Need”).

### What we are not claiming

Not all immune mechanisms will translate to AI. The convergences concern recognition and generalization, not the full complexity of either system. Other immune processes—complement activation, cytokine signaling, innate immunity—may have no meaningful AI parallel. Other AI architectures—convolutional networks, diffusion models—may have no immune parallel. The framework addresses a specific computational problem, not a universal correspondence between fields.

### Selection pressure

The selection pressure on adaptive immunity is singular in biology (*79*). An organism that fails to recognize a novel pathogen dies. An organism that fails to discriminate self from non-self develops autoimmunity. There is no partial success, no compensatory mechanism. This pressure has operated continuously for 500 million years (*80*), across all jawed vertebrates, in every environment pathogens occupy. The computational strategies that emerged—Boltzmann-weighted recognition, contrastive selection, hierarchical optimization, retrieval-augmented memory—were optimized for survival across half a billion years of pathogen co-evolution.

### Adaptive immunity as intelligence

If intelligence is defined as the capacity for recognition and generalization under resource constraints—learning to respond appropriately to novel situations based on limited prior experience—then the adaptive immune system is a form of intelligence. Not metaphorically, but computationally. It recognizes virtually any molecular structure (*22*), discriminates self from non-self across an antigen space exceeding 10^15^ possibilities (*23*), and forms memories lasting decades (*81*). It accomplishes this with 10^7^–10^9^ lymphocytes - a compression that no artificial system has matched (*20, 21*).

### Implications for immune specificity prediction

If transformer attention is mathematically equivalent to antibody-antigen binding, then transformers are the natural architecture for modeling immune recognition—the tool matches the problem by shared mathematical structure. This suggests that accurate sequence-to-specificity prediction—determining which antibodies bind which antigens from sequence alone—is not only possible but inevitable at repertoire-proteome-scale, given sufficient training data. The problem is learnable because evolution learned it. The architecture is correct because it implements the same computation. What remains is data (*82*).

### The convergence

Modern AI arrived at these strategies through gradient descent on benchmark performance (*83*). The softmax function emerged as a useful nonlinearity; it is also the maximum entropy distribution for converting energies to probabilities (*84*). InfoNCE emerged as an effective contrastive objective; it is also the mathematics of selection (*2*). The training hierarchy emerged as best practice; it is also the structure of affinity maturation (*54*). These were empirical discoveries in AI. They are mathematical necessities in immunology.

When two independent optimization processes—one biological, one computational—arrive at identical solutions, those solutions likely reflect the structure of the problem itself (*85*). Evolution solved the recognition-generalization problem 500 million years ago. We are only now understanding the solution.

## Acknowledgments

Thanks to Thomas Bikias and Mason Minot for reading the manuscript and providing feedback.

## Author contributions

S.T.R. conceived the study, developed the theoretical framework, and wrote the manuscript.

## Competing interests

S.T.R. is a co-founder and scientific advisor of Engimmune Therapeutics, Encelta, Fy Cappa Biologics and BIIE.AI.

## Data and materials availability

This is a theoretical paper. No new data was generated.

## Supplementary Materials

### Supplementary Note S1 Deriving Transformer Attention from Antibody-Antigen Binding

#### Purpose

This note provides the complete mathematical derivation of the transformer attention mechanism from the biophysics of antibody-antigen binding. Each step is explicit to enable verification.

##### S1.1 Notation Summary

| Symbol | Definition | Domain |
| --- | --- | --- |
| $i$ | Paratope residue index (antibody) | $i \in 1, \dots, n_P$ |
| $j$ | Epitope residue index (antigen) | $j \in 1, \dots, n_E$ |
| $E_{ij}$ | Interaction energy between residues $i$ and $j$ | $E_{ij} \in \mathbb{R}$ (negative = favorable) |
| $k_B$ | Boltzmann constant | $1.38 \times 10^{-23}$ J/K |
| $T$ | Temperature | $T > 0$ (Kelvin) |
| $s_{ij}$ | Dimensionless compatibility score | $s_{ij} = -E_{ij}/k_B T$ |
| $P(j\vert i)$ | Probability residue $i$ contacts residue $j$ | $P(j\vert i) \in [0, 1], \sum_j P(j\vert i) = 1$ |
| $\mathbf{q}_i$ | Query vector for position $i$ | $\mathbf{q}_i \in \mathbb{R}^{d_k}$ |
| $\mathbf{k}_j$ | Key vector for position $j$ | $\mathbf{k}_j \in \mathbb{R}^{d_k}$ |
| $\mathbf{v}_j$ | Value vector for position $j$ | $\mathbf{v}_j \in \mathbb{R}^{d_v}$ |
| $A_{ij}$ | Attention weight from $i$ to $j$ | $A_{ij} \in [0, 1], \sum_j A_{ij} = 1$ |

##### S1.2 Immunological Setup

###### Physical system

An antibody (Ab) binds an antigen (Ag) through contact between the paratope (Ab binding surface) and epitope (Ag target region).

###### Contact model

Each paratope residue may contact each epitope residue *j*. The contact probability depends on the interaction energy *E*_*ij*_, which captures:

- Electrostatic interactions
- Hydrophobic contacts
- Hydrogen bonds
- Van der Waals forces
- Shape complementarity

###### Assumption

The system is at thermal equilibrium. This is standard for analyzing molecular binding.

##### S1.3 Boltzmann Distribution Derivation

###### Statement

At thermal equilibrium, the probability that paratope residue *i* contacts epitope residue *j* is:

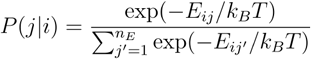

###### Derivation

*Step 1:* By the fundamental postulate of statistical mechanics, the probability of a microstate with energy *E* is proportional to exp (− *E*_*ij*_ /*k*_*B*_ *T*) .

*Step 2:* For residue *i* choosing among possible contacts *j* ∈ 1, …, *n*_*E*_, each contact has energy *E*_*ij*_.

*Step 3:* The unnormalized probability of contact *j* is exp (− *E*_*ij*_ /*k*_*B*_ *T*) .

*Step 4:* Normalizing over all possible contacts:

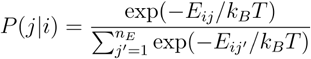

###### Verification

- *P*(*j*|*i*) ≥ 0for all *j* ✓ (exponential is always positive)
- 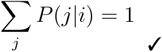 (by construction of denominator)

Lower energy → higher probability ✓ (more negative *E*_*ij*_ → larger numerator)

##### S1.4 Change of Variables: Energy to Compatibility Score

###### Definition

Define the dimensionless compatibility score:

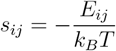

###### Interpretation

- *s*_*ij*_ > 0 when *s*_*ij*_ < 0 (favorable interaction)
- *s*_*ij*_ < when *E*_*ij*_ > (unfavorable interaction)
- Higher *s*_*ij*_ = better compatibility

###### Substitution

Replace − *E*_*ij*_ /*k*_*B*_ *T* with : *s*_*ij*_ :

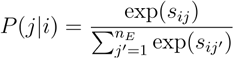

This is the **softmax function** applied to scores 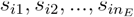.

##### S1.5 The Softmax-Boltzmann Identity

###### Claim

The softmax function and Boltzmann distribution are mathematically identical.

###### Softmax definition

For a vector S=(*s*_1_,…,*s*_*n*_):

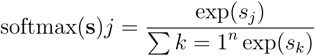

###### Boltzmann (in terms of compatibility scores)

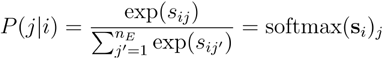

where 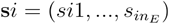.

###### Conclusion

The probability distribution over contacts IS a softmax over compatibility scores. This is not an approximation - it is an identity.

##### S1.6 Query-Key Parameterization

###### Motivation

How should compatibility scores *s*_*ij*_ be computed?

###### Biophysical intuition

Compatibility depends on:

- Features of the paratope residue *i* (charge, size, hydrophobicity, shape)
- Features of the epitope residue *j* (charge, size, hydrophobicity, shape)
- How well these features complement each other

###### Parameterization

Represent residue features as vectors:

- Paratope residue *i* : feature vector **P**_*i*_ ∈ ℝ^*d*^
- Epitope residue *j*: feature vector **e**_*i*_ ∈ ℝ^*d*^

###### Compatibility as dot product

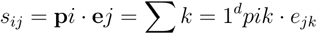

###### Justification

The dot product measures alignment between feature vectors:

- Parallel vectors (similar features) → high dot product → high compatibility
- Orthogonal vectors (unrelated features) → zero dot product → neutral
- Anti-parallel vectors (opposing features) → negative dot product → low compatibility

###### Renaming for transformer notation

- Paratope features → Query: **q**_*i*_ *=* **P**_*i*_
- Epitope features → Key: k_*j*_ *=* **e**_*j*_

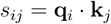

##### S1.7 Scaling Factor

###### Issue

For high-dimensional vectors, dot products can have large magnitude, making softmax outputs very sharp.

###### In physics

Temperature *T* controls distribution sharpness. Low *T* → sharp; high *T* → diffuse.

###### In transformers

Use scaling factor 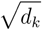 where *d*_*k*_ is the key dimension:

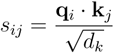

###### Correspondence

- Boltzmann: *P* ∝ exp (− *E*_*ij*_ */k*_*B*_ *T*)
- Attention: 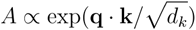

The scaling factor 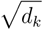 plays the role of temperature.

##### S1.8 Value Aggregation

###### Question

What happens after contact probabilities are computed?

###### Biophysical answer

When residue *i* contacts residue *j*, properties of *j* influence the state of *i*. The total influence on *i* is a weighted sum over all possible contacts:

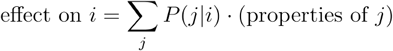

###### Parameterization

Represent the information transferred by residue *j* as a value vector **v**_*j*_ :

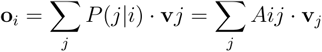

This is the attention output - a weighted average of value vectors, where weights are contact probabilities (attention weights).

##### S1.9 Complete Attention Equation

###### Assembly

Combining all components:

*Step 1:* Compute compatibility scores:

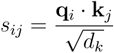

*Step 2:* Compute attention weights via softmax:

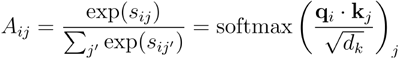

*Step 3:* Compute output as weighted sum of values:

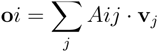

###### Matrix form

For queries *Q*, keys *K*, values *V*:

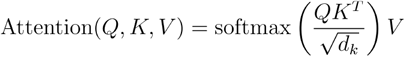

**This is the transformer attention equation (Vaswani et al., 2017)**

##### S1.10 Multi-Head Attention from CDR Loops

###### Observation

Antibodies have 6 CDR loops (CDRH1, CDRH2, CDRH3, CDRL1, CDRL2, CDRL3), each engaging different epitope regions.

###### Correspondence

Multi-head attention uses *H* parallel attention mechanisms:

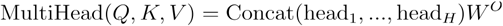

where each head has independent parameters:

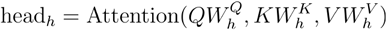

###### Mapping

**Mapping:**
| CDR Loops | Multi-Head Attention |
| --- | --- |
| 6 CDR loops | $H$ attention heads |
| Each loop has distinct sequence/structure | Each head has distinct $W_h^Q, W_h^K, W_h^V$ |
| Different loops contact different epitope regions | Different heads attend to different patterns |
| Loops contribute different interaction types | Heads specialize for different relationships |
| Combined paratope = union of CDR contributions | Combined output = concatenation of head outputs |

##### S1.11 Summary of Derivation

| Step | Starting Point | Result |
| --- | --- | --- |
| 1 | Thermal equilibrium | Boltzmann distribution over contacts |
| 2 | Change of variables | $P(j\vert i) = \text{softmax}(s_i)_j$ |
| 3 | Feature representation | $s_{ij} = \mathbf{q}_i \cdot \mathbf{k}_j$ |
| 4 | Temperature/scaling | $s_{ij} = \mathbf{q}_i \cdot \mathbf{k}_j / \sqrt{d_k}$ |
| 5 | Information transfer | $\mathbf{o}_i = \sum_j A_{ij} \mathbf{v}_j$ |
| 6 | Parallel channels | Multi-head attention |

###### Final result

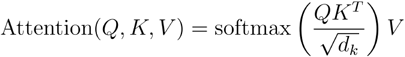

##### S1.12 Verification Checklist

- [ ] Boltzmann distribution formula is correct
- [ ] Change of variables preserves probability properties
- [ ] Softmax-Boltzmann equivalence is exact (not approximate)
- [ ] Dot product parameterization is justified
- [ ] Scaling factor corresponds to temperature
- [ ] Value aggregation follows from contact-weighted information transfer
- [ ] Multi-head attention maps to CDR loop structure
- [ ] Final equation matches Vaswani et al. (2017)

### Supplementary Note S2 Deriving Contrastive Learning from Clonal Selection

#### Purpose

This note provides the complete mathematical derivation of the InfoNCE contrastive learning objective from thymic and germinal center selection dynamics.

##### S2.1 Notation Summary

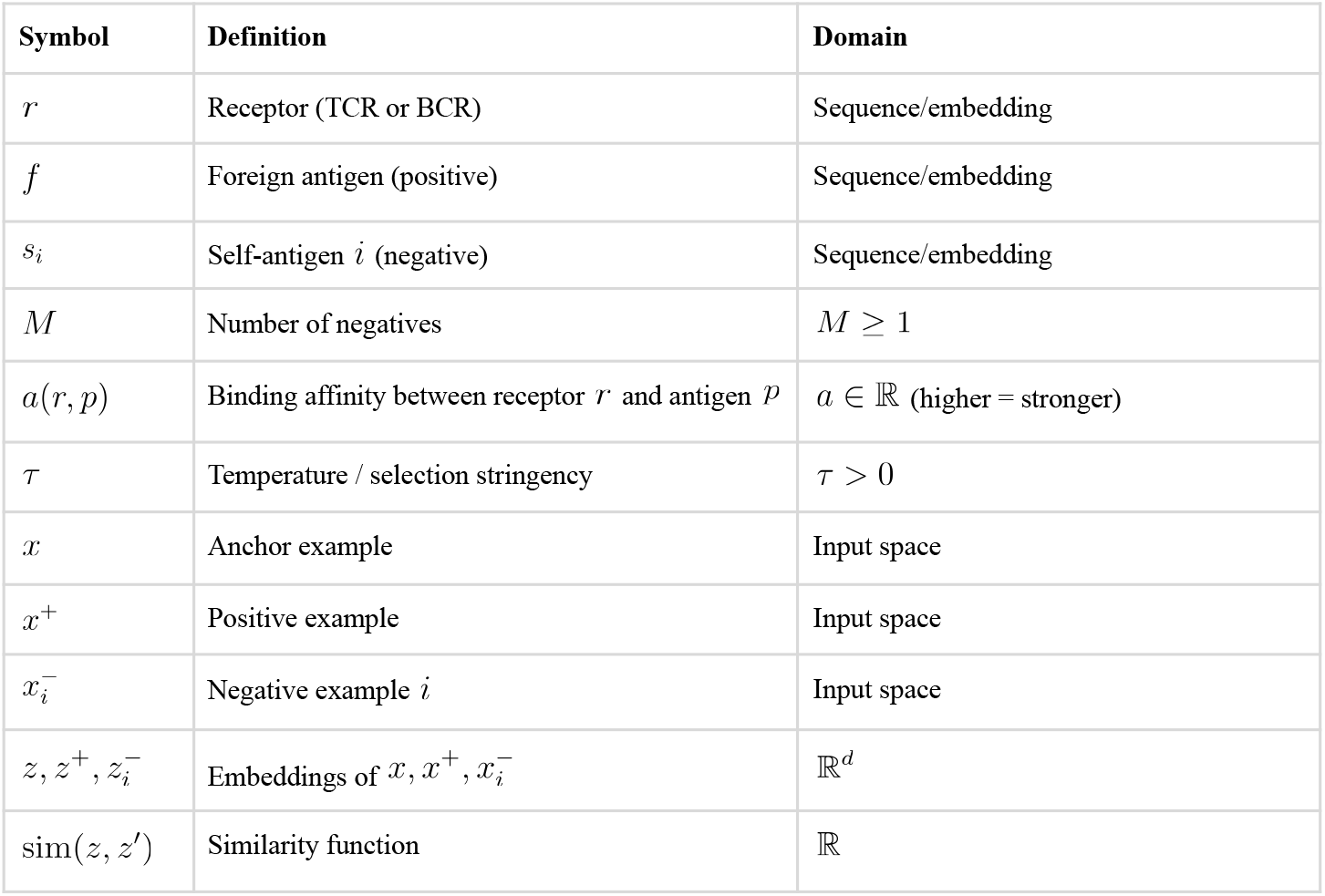

| Symbol | Definition | Domain |
| --- | --- | --- |
| $r$ | Receptor (TCR or BCR) | Sequence/embedding |
| $f$ | Foreign antigen (positive) | Sequence/embedding |
| $s_i$ | Self-antigen $i$ (negative) | Sequence/embedding |
| $M$ | Number of negatives | $M \geq 1$ |
| $a(r, p)$ | Binding affinity between receptor $r$ and antigen $p$ | $a \in \mathbb{R}$ (higher = stronger) |
| $\tau$ | Temperature / selection stringency | $\tau > 0$ |
| $x$ | Anchor example | Input space |
| $x^+$ | Positive example | Input space |
| $x_i^-$ | Negative example $i$ | Input space |
| $z, z^+, z_i^-$ | Embeddings of $x, x^+, x_i^-$ | $\mathbb{R}^d$ |
| $\text{sim}(z, z')$ | Similarity function | $\mathbb{R}$ |

##### S2.2 Immunological Setup: Thymic Selection

###### Physical system

T cell development in the thymus.

###### Process

1. Developing T cells (thymocytes) express T cell receptors (TCRs) generated by V(D)J recombination
2. Each TCR is tested against peptides presented on MHC molecules
3. **Positive selection:** TCRs must bind self-MHC with sufficient affinity to survive
4. **Negative selection:** TCRs that bind self-peptide-MHC too strongly are eliminated

###### Goal

Produce a repertoire that:

- Responds to foreign peptides presented on self-MHC
- Tolerates self-peptides (no autoimmunity)

This is a **contrastive discrimination task**: select the positive (foreign) while rejecting negatives (self).

##### S2.3 Selection Probability Derivation

###### Setup

A receptor encounters:

- One foreign antigen *f* (positive)
- *M* self-antigens *s*_1_,…, *s*_*M*_ (negatives)

###### Assumption

Selection follows competitive dynamics—cells compete for survival signals based on binding affinity.

###### Model

Under Boltzmann-like competition, the probability of selecting antigen *j* from a candidate set is proportional to exp (*a*(*r,j*)/ *τ* ), where *a*(*r,j*) is the affinity and *τ* controls stringency.

###### Derivation

*Step 1:* Unnormalized selection weight for foreign antigen: *w*_*f*_ *=* exp (*a* ( *r,f*)/ *τ*)

*Step 2:* Unnormalized selection weight for each self-antigen: 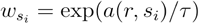

*Step 3:* Total unnormalized weight:

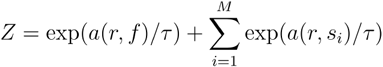

*Step 4:* Probability of selecting foreign antigen:

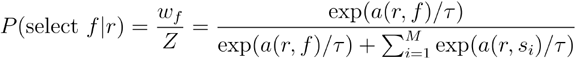

###### Interpretation

This is the probability that a receptor *r* survives selection for the foreign antigen against a background of self-antigens.

##### S2.4 Contrastive Learning Setup

###### Setup

An anchor *x* is paired with:

- One positive *x*^+^(similar to anchor)
- *N* negatives 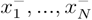 (dissimilar to anchor)

###### Embeddings

An encoder *f* _*θ*_ maps inputs to embeddings:

- *z*= *f* _*θ*_ (*x*)
- *z*^+^ = *f* _*θ*_ (*x*)
- 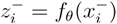

###### Similarity

Typically cosine similarity or dot product:

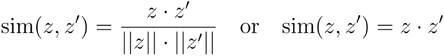

###### Goal

Learn embeddings such that:

- sim (*z, z*^+^ ) is high (positives are close)
- 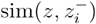 is low (negatives are far)

##### S2.5 The Mathematical Equivalence

###### Contrastive selection probability

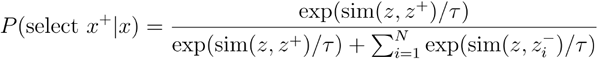

###### Immune selection probability

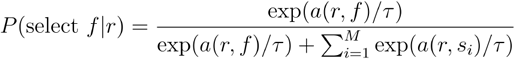

###### Correspondence

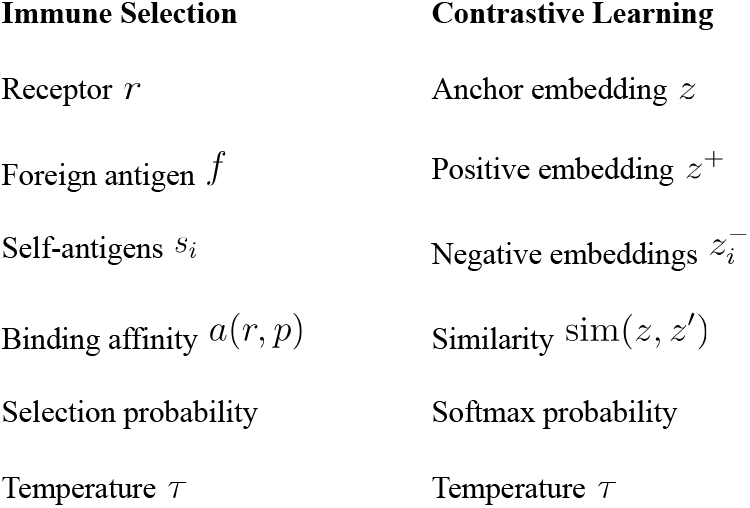

###### The equations are identical

This is not an analogy—it is the same mathematical structure.

##### S2.6 InfoNCE as Negative Log-Likelihood

###### Definition

The InfoNCE loss is the negative log of selection probability:

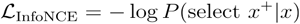

###### Expanded form

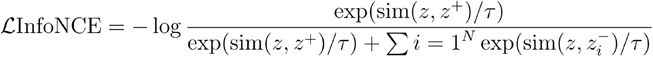

###### Simplified

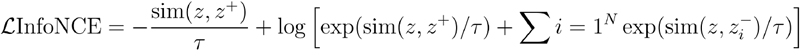

###### Interpretation

Minimizing InfoNCE maximizes the probability of correctly selecting the positive from among the negatives.

###### Immunological interpretation

This is exactly the training objective implicit in thymic selection—maximize the probability that receptors select foreign antigens over self-antigens.

##### S2.7 In-Batch Negatives

###### Common practice

Use other examples in the batch as negatives.

For a batch of *B* examples 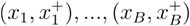:

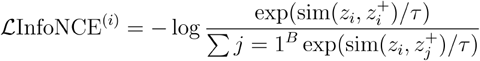

###### Immunological interpretation

This corresponds to selection among a cohort of developing lymphocytes, where each cell’s positive (foreign antigen) competes against other cells’ positives (which serve as “self” for the focal cell).

##### S2.8 Hard Negatives and AIRE

###### Contrastive learning insight

Hard negatives—negatives similar to the positive—improve discrimination. Easy negatives provide little gradient signal.

###### Immunological implementation: AIRE

AIRE (Autoimmune Regulator) is a transcription factor expressed in thymic medullary epithelial cells.

###### Function

AIRE drives ectopic expression of tissue-restricted antigens (TRAs)—proteins normally expressed only in specific organs:

- Insulin (pancreas)
- Thyroglobulin (thyroid)
- Myelin basic protein (nervous system)

**Why express pancreatic proteins in the thymus?** These are **hard negatives**

###### Mechanism

1. Without AIRE: T cells only see ubiquitous self-antigens
2. T cells that cross-react with TRAs escape negative selection
3. Upon peripheral exposure to TRAs, these T cells attack self-tissue → autoimmunity

###### With AIRE

1. TRAs are presented in thymus
2. T cells cross-reactive with TRAs are deleted
3. Tolerance extends to tissue-restricted self

###### Correspondence

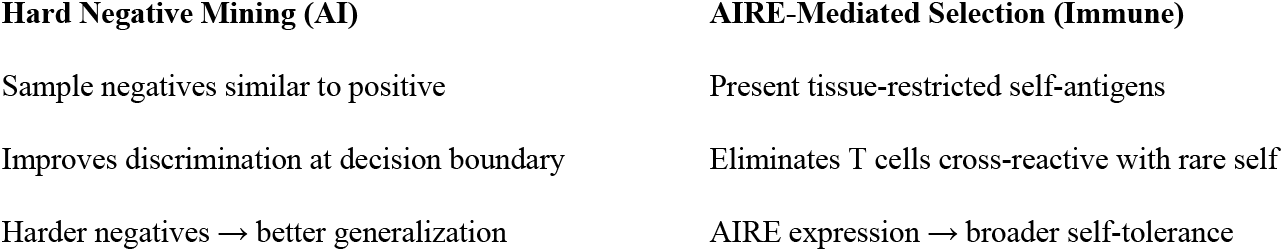

###### Clinical evidence

AIRE deficiency causes Autoimmune Polyendocrine Syndrome Type 1 (APS-1):

- Autoimmunity against multiple organs
- Biological equivalent of training without hard negatives

##### S2.9 Temperature Parameter

###### Mathematical role

Temperature *τ* controls distribution sharpness.

###### Low *τ* (stringent selection)

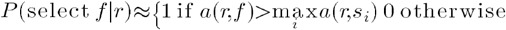

- Small affinity differences → large probability differences
- Only highest-affinity interaction dominates
- Sharp discrimination

###### High *τ* (permissive selection)

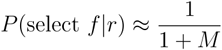

- Probabilities approach uniform
- Multiple candidates have similar probability
- Diffuse discrimination

###### Immunological interpretation

- Low *τ*: Stringent selection → highly specific repertoire, but may delete useful receptors
- High *τ*: Permissive selection → diverse repertoire, but may retain autoreactive receptors

###### AI interpretation

- Low *τ*: Sharp embeddings, may overfit to training negatives
- High : *τ* Smooth embeddings, may fail to discriminate

###### The same parameter controls the same tradeoff

##### S2.10 Germinal Center Selection

The same mathematics applies to B cell selection in germinal centers.

###### Setup

B cells compete for T follicular helper (Tfh) cell help based on ability to capture and present antigen.

###### Process

1. B cells capture antigen via surface BCR
2. Higher-affinity B cells capture more antigen
3. B cells present antigen-derived peptides on MHC class II
4. Tfh cells provide help to B cells presenting cognate peptide
5. B cells receiving more help proliferate; others die

###### Mathematical form

For *B*_*GC*_ competing B cells with affinities 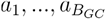:

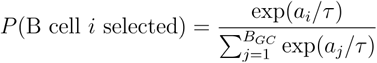

This is softmax competition - the same equation as contrastive learning batches.

##### S2.11 Summary of Derivation

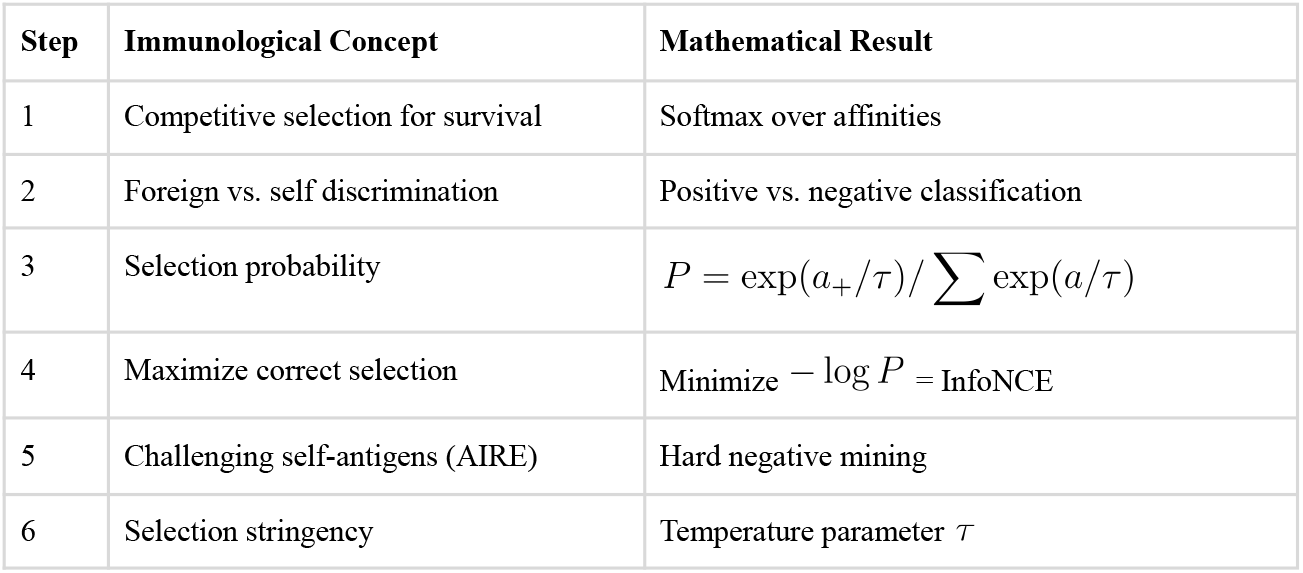

| Step | Immunological Concept | Mathematical Result |
| --- | --- | --- |
| 1 | Competitive selection for survival | Softmax over affinities |
| 2 | Foreign vs. self discrimination | Positive vs. negative classification |
| 3 | Selection probability | $P = \exp(a_+/\tau) / \sum \exp(a/\tau)$ |
| 4 | Maximize correct selection | Minimize $-\log P = \text{InfoNCE}$ |
| 5 | Challenging self-antigens (AIRE) | Hard negative mining |
| 6 | Selection stringency | Temperature parameter $\tau$ |

###### Final result

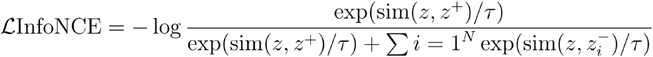

This is derivable from:

- Clonal selection theory (Burnet, 1959)
- Thymic selection mechanisms (established 1980s)
- AIRE function (Anderson et al., 2002)

No machine learning knowledge required.

##### S2.12 Verification Checklist

- [ ] Selection probability formula is correct softmax form
- [ ] InfoNCE equals negative log of selection probability
- [ ] Temperature interpretation matches Boltzmann physics
- [ ] AIRE function correctly described
- [ ] Germinal center selection follows same mathematics
- [ ] Correspondence table entries are accurate

### Supplementary Note S3 Training Hierarchy from Affinity Maturation

#### Purpose

This note formalizes the correspondence between the three-stage AI training hierarchy (pre-training → fine-tuning → RLHF) and antibody affinity maturation (germline → SHM → Tfh selection).

##### S3.1 Notation Conventions

To avoid symbol collision across stages, we use:

###### Immune system

Immune system:
| Symbol | Definition |
| --- | --- |
| $s$ | Antibody sequence |
| $\mathcal{S}$ | Space of all antibody sequences |
| $\mathcal{V}, \mathcal{D}, \mathcal{J}$ | V, D, J gene segment sets |
| $P_{\text{germline}}(s)$ | Germline distribution over sequences |
| $\mu$ | SHM mutation rate (per bp per division) |
| $q_i$ | Quality score for B cell $i$ (Tfh selection) |

###### AI system

AI system:
| Symbol | Definition |
| --- | --- |
| $\theta$ | Model parameters |
| $\theta_0$ | Pre-trained parameters |
| $\Theta$ | Parameter space |
| $P_{\theta}(y\vert x)$ | Model's conditional distribution |
| $\eta$ | Learning rate |
| $R_i$ | Reward score for response $i$ (RLHF) |

##### S3.2 Stage 1: Germline as Optimized Prior

###### S3.2.1 Germline Generative Process

Antibodies are assembled from gene segments via V(D)J recombination:

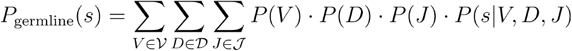

where:

- *P* (*V*), *P* (*D*), *P* (*J* ) are segment usage frequencies (shaped by evolution)
- *P* (*s* | *V, D, J*) captures junctional diversity from random insertions, deletions, and N-nucleotides

###### S3.2.2 Properties of Germline Distribution

The distribution *P* _germline_ (*s*)is **non-uniform**:

- Some sequences are much more probable than others
- Evolution has biased toward sequences that:
  ∘ Fold into stable antibody structures
  ∘ Bind common pathogen motifs (e.g., VH1-69 for influenza)
  ∘ Provide useful combinatorial diversity

###### Key insight

The germline encodes an **optimized prior**—a starting distribution much better than random.

###### S3.2.3 Pre-trained Weights as Learned Distribution

A pre-trained language model with parameters *θ*_0_ defines:

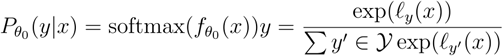

where 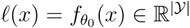 are logits over vocabulary 𝒴.

###### Properties

This distribution reflects patterns in pre-training data:

- Grammar and syntax
- World knowledge
- Reasoning patterns
- Common response structures

###### S3.2.4 Structural Correspondence

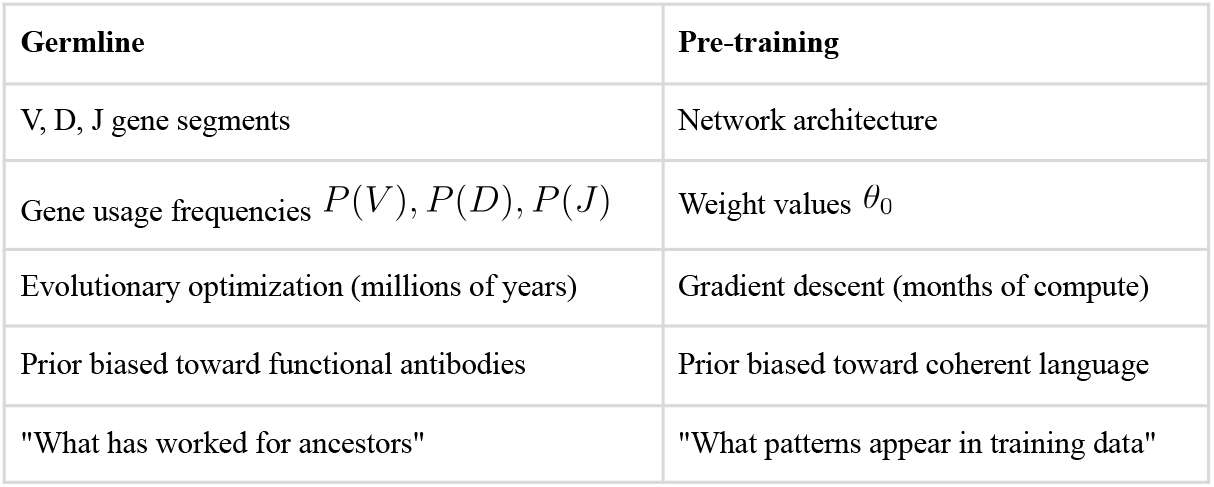

| Germline | Pre-training |
| --- | --- |
| V, D, J gene segments | Network architecture |
| Gene usage frequencies $P(V), P(D), P(J)$ | Weight values $\theta_0$ |
| Evolutionary optimization (millions of years) | Gradient descent (months of compute) |
| Prior biased toward functional antibodies | Prior biased toward coherent language |
| "What has worked for ancestors" | "What patterns appear in training data" |

**Both encode starting points much better than random.**

##### S3.3 Stage 2: SHM as Stochastic Local Search

###### S3.3.1 Somatic Hypermutation Dynamics

When a B cell enters a germinal center, the enzyme AID introduces point mutations.

###### Mutation operator

Let *s* _*t*_ denote the antibody sequence at round *t*:

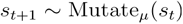

where Mutate_*μ*_ is a stochastic operator introducing mutations at rate *μ*.

###### Rate

*μ* ≈10^−3^ mutations per base pair per cell division.

###### Expected mutations per round

For variable region length *L* ≈ 1400 bp:

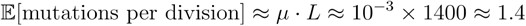

###### Targeting

Mutations concentrate in CDRs (binding regions) while relatively sparing framework regions. AID preferentially targets certain sequence motifs (WRC/GYW hotspots).

###### S3.3.2 Fine-tuning Dynamics

Fine-tuning adapts pre-trained *θ*_0_ weights to a specific task:

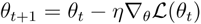

where:

- *η* ≈ 10^−3^ to 10^−5^ is the learning rate
- ℒ( *θ* ) is the task-specific loss

###### Layer-wise rates

Common practice uses different learning rates for different layers:

- Higher rates for task-specific (later) layers
- Lower or frozen rates for general (early) layers

This parallels CDR vs. framework targeting in SHM.

###### S3.3.3 Critical Distinction

The SHM operator *s*_*t+*1_ ∼ Mutate *μ* (*s*_*t*_) *and gradient update θt* + 1 = *θ*_*t*_ − *η* ∇ ℒ are **not mathematically identical**:

- SHM: Stochastic edit on discrete sequences
- Fine-tuning: Directed step in continuous parameter space

However, SHM + selection effectively creates **directed optimization**: random variation plus selection approximates gradient-free optimization.

###### S3.3.4 Structural Correspondence

| SHM | Fine-tuning |
| --- | --- |
| Mutation rate $\mu \approx 10^{-3}$ per bp | Learning rate $\eta \approx 10^{-3}$ to $10^{-5}$ |
| Stochastic point mutations | Gradient-directed updates |
| Concentrated in CDRs | Can be concentrated in task-specific layers |
| Multiple rounds over ~3-4 weeks | Multiple epochs over hours to days |
| Starting from germline sequence | Starting from pre-trained weights |

###### Key similarity

Both use step sizes on the order of 10^−3^ . This may reflect a shared constraint—the “Goldilocks zone” for adaptation from an optimized prior. Too large → destroy prior value. Too small → insufficient adaptation.

##### S3.4 Stage 3: Tfh Selection as Preference Feedback

###### S3.4.1 T Follicular Helper Selection

After SHM generates variants, which survive? T follicular helper (Tfh) cells determine this.

###### Process

1. B cell captures antigen via BCR
2. B cell processes and presents antigen-derived peptides on MHC class II
3. Tfh cells recognize peptide-MHC and provide help signals (CD40L, cytokines)
4. B cells receiving more help proliferate; those receiving less die

###### Critical observation

Tfh cells do not measure antibody-antigen affinity directly. They measure **antigen presentation quality** - a proxy that correlates with affinity but is not identical.

The B cell receives feedback: “you’re doing well” (survival signal) or “you’re not doing well” (death), not “your affinity is X nM.”

###### S3.4.2 Selection Mathematics

For a set of competing B cells 𝒞with quality scores 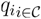:

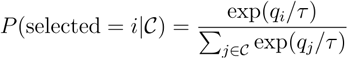

This is softmax competition—the same form as Convergences 1 and 2.

###### S3.4.3 RLHF Process

RLHF (Reinforcement Learning from Human Feedback) aligns model behavior with human preferences:

1. Model generates multiple candidate responses to prompts
2. Human raters compare responses: “A is better than B”
3. A reward model *R* is trained to predict human preferences
4. The language model is optimized toward higher reward

###### Critical observation

Humans do not specify the correct answer. They provide **preference feedback**—a signal that response A is better than B.

###### S3.4.4 RLHF Mathematics

For responses with reward scores 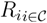:

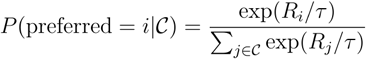

###### S3.4.5 Mathematical Equivalence

Stage 3 exhibits **exact mathematical equivalence**:

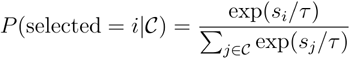

where:

- In Tfh selection: *s*_*i*_ = *q*_*i*_ (quality proxy from antigen presentation)
- In RLHF: *s*_*i*_ = *R*_*i*_ (reward model score from human preferences)

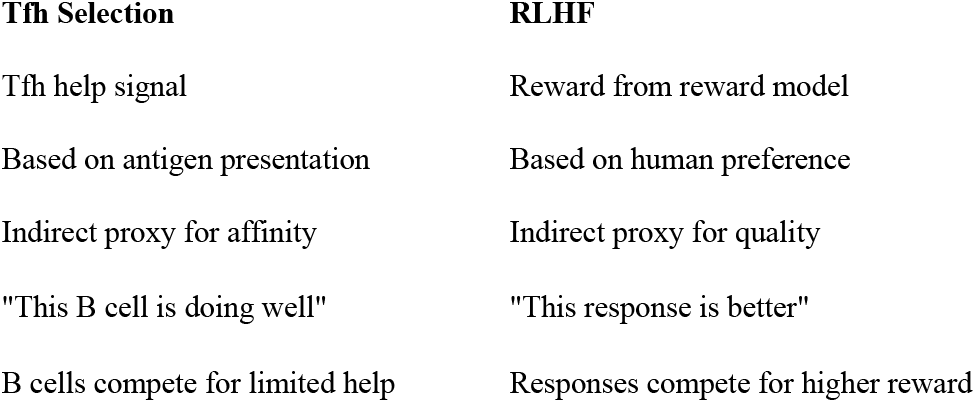

##### S3.5 Why Three Stages?

###### S3.5.1 Different Objectives Require Different Feedback

| Stage | Objective | Data Source | Feedback Type | Characteristic |
| --- | --- | --- | --- | --- |
| 1 | General capability | Evolutionary history / corpus | Implicit (survival / prediction) | Expensive to create, reusable |
| 2 | Task-specific performance | Current antigen / task data | Explicit (affinity / task loss) | Efficient from good starting point |
| 3 | Response calibration | Tfh interaction / human preference | Comparative preference | Expensive per example, limited scale |

###### One stage fails because

- Cannot efficiently combine general learning with task-specific optimization
- Cannot do RLHF-style training with trillions of examples (too expensive)
- Conflicting objectives create tension

###### Two stages (no Stage 3) fails because

- Capability optimization doesn’t guarantee alignment
- A highly capable model/antibody can still:
  ∘ Generate harmful outputs / cause immunopathology
  ∘ Have wrong tone, style, or magnitude
  ∘ Violate user/host expectations
- Stage 3 provides calibration that capability optimization alone cannot achieve

###### Both systems embody

**Invest heavily in general capability because it amortizes across all future specific challenges**.

###### Germline transfer

The same germline V gene (e.g., VH1-69) contributes to antibodies against influenza, HIV, hepatitis C, and SARS-CoV-2. Evolution optimized germline genes to be useful across many antigens.

###### Pre-training transfer

The same pre-trained weights enable fine-tuning for summarization, translation, coding, question-answering. Companies invest massively in pre-training because it enables future specialization at low marginal cost.

##### S3.6 Summary Table

| Stage | Immune System | AI Training | Math Status |
| --- | --- | --- | --- |
| 1 | $P_{\text{germline}}(s)$ | $P_{\theta_0}(y\vert x)$ | Structural correspondence |
| 2 | $s_{t+1} \sim \text{Mutate}_{\mu}(s_t)$ | $\theta_{t+1} = \theta_t - \eta \nabla \mathcal{L}$ | Structural correspondence |
| 3 | $P \propto \exp(q_i/\tau)$ | $P \propto \exp(R_i/\tau)$ | **Exact equivalence** |

The convergence is **strategic** for Stages 1-2 (same structure, analogous roles) and **exact** for Stage 3 (identical softmax selection mathematics).

##### S3.7 Verification Checklist

- [ ] Germline distribution formula correctly represents V(D)J recombination
- [ ] SHM mutation rate ( *μ* = 10^−3^ ) is correct
- [ ] Fine-tuning learning rate ( *η* = 10^−3^) is typical
- [ ] Tfh selection follows softmax competition
- [ ] RLHF reward optimization follows same softmax form
- [ ] Stage 3 equations are mathematically identical
- [ ] Three-stage necessity arguments are logically sound

### Supplementary Note S4 Dual Memory Architecture from Immune Memory

#### Purpose

This note formalizes the correspondence between Retrieval-Augmented Generation (RAG) and the immune dual memory system (plasma cells + memory B cells).

##### S41.1 Notation Summary

###### Immune system

Immune system:
| Symbol | Definition |
| --- | --- |
| $a$ | Antigen (query) |
| $\mathcal{B}$ | Memory B cell pool |
| $i, j$ | B cell indices |
| $A_i(a)$ | Affinity of B cell $i$'s BCR for antigen $a$ |
| $\tau$ | Activation stringency |
| LLPC | Long-lived plasma cell |
| $f_p$ | Encounter frequency for pathogen $p$ |
| $s_p$ | Severity of pathogen $p$ |

###### RAG system

RAG system:
| Symbol | Definition |
| --- | --- |
| $q$ | Query |
| $\mathcal{D}$ | Document store |
| $d_j$ | Document $j$ |
| $S(q, d)$ | Similarity between query $q$ and document $d$ |
| $k$ | Number of documents to retrieve |

##### S4.2 Immune Dual Memory Architecture

###### Location

Bone marrow survival niches

###### Properties

- Constitutively secrete antibody without requiring reactivation
- Provide immediate, always-on protection
- Limited capacity: ∼10^4^ − 10^5^ niches available
- Metabolically expensive (continuous protein synthesis)
- Long-lived (decades in humans)
- Difficult to update (require niche competition to replace)

###### Function

Maintain baseline antibody titers against previously encountered pathogens.

###### Location

Circulate in blood and lymphoid tissues

###### Properties

- Quiescent until antigen re-encounter
- Require activation to produce antibody
- Large capacity: ∼10^7^ – 10^8^ cells
- Metabolically inexpensive when dormant
- Long-lived but require periodic antigen exposure
- Easy to expand upon re-challenge

###### Function

Rapid, amplified secondary response upon re-infection.

###### S4.2.3 Dual System Properties

| Property | LLPCs | Memory B Cells |
| --- | --- | --- |
| Activation | Constitutive | On-demand |
| Response latency | Immediate | Delayed (days) |
| Capacity | Limited | Large |
| Metabolic cost | High | Low (dormant) |
| Update cost | High | Low |
| Flexibility | Fixed specificity | Can undergo further affinity maturation |

##### S4.3 RAG Dual Memory Architecture

###### Location

Neural network parameters *θ*

###### Properties

- Knowledge encoded in weight values
- Always available during generation
- Provides immediate, context-free responses
- Limited by model capacity
- Expensive to update (requires retraining)
- Cannot easily add new knowledge

###### Location

External searchable database 𝒟

###### Properties

- Documents stored as text + embeddings
- Retrieved on-demand based on query similarity
- Requires retrieval step before use
- Virtually unlimited capacity
- Cheap to update (add/remove documents)
- Provides fresh, specific information

###### S4.3.3 Dual System Properties

| Property | Parametric Memory | Retrieved Memory |
| --- | --- | --- |
| Activation | Always available | On-demand |
| Response latency | Immediate | Requires retrieval |
| Capacity | Limited (model size) | Virtually unlimited |
| Compute cost | High (large model) | Low (storage only) |
| Update cost | High (retraining) | Low (add documents) |
| Flexibility | Fixed until retrained | Can add new knowledge |

##### S4.4 Structural Correspondence

| Immune Memory | RAG |
| --- | --- |
| Long-lived plasma cells | Parametric memory (weights) |
| Memory B cells | Document store |
| Constitutive antibody secretion | Always-available model outputs |
| Antigen-triggered reactivation | Query-triggered retrieval |
| BCR affinity for antigen | Embedding similarity for query |
| Limited bone marrow niches | Limited model capacity |
| Large memory B cell pool | Large document database |

##### S4.5 Similarity-Based Retrieval

###### S4.5.1 Memory B Cell Activation

When antigen *a* is encountered, memory B cells compete for activation based on BCR affinity.

###### Probabilistic activation

For memory B cell pool ℬ *=* 1, … *N*:

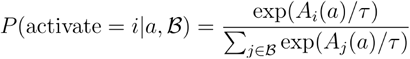

where:

- *A*_*i*_ (*a* ) is affinity of cell *i*’s BCR for antigen *a*
- *τ* controls activation stringency

###### Interpretation

Higher-affinity cells are more likely to be activated.

###### S4.5.2 Document Retrieval

When query *q* is presented, documents compete for retrieval based on similarity.

###### Probabilistic retrieval

For document store 𝒟= *d*_1,_ …, *d*_*M*_:

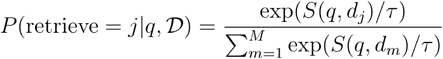

where:

- *S* (*q, d*_*j*_ ) is similarity between query and document (e.g., cosine similarity of embeddings)
- *τ* controls retrieval sharpness

###### S4.5.3 Mathematical Equivalence

Both mechanisms share identical structure:

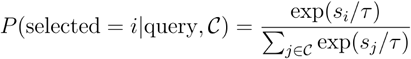

where:

- In immune memory: *s*_*i*_ = *A*_*i*_ (*a*) (BCR affinity)
- In RAG: *s*_*i*_ *= S* (*q, d*_*i*_ ) (embedding similarity)

This is the same Boltzmann-form softmax appearing in Convergences 1, 2, and 3.

###### S4.6.1 Cosine Similarity (RAG)

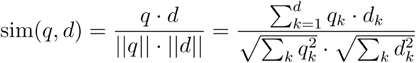

###### Properties

- Dimensionless, range [ℒ 1, 1]
- 1 = identical direction
- 0 = orthogonal
- -1 = opposite direction

###### S4.6.2 BCR-Antigen Affinity (Immune)

BCR-antigen binding follows the same biophysics as antibody-antigen binding (Convergence 1):

- Governed by Boltzmann distribution over contact energies
- Affinity = − Δ *G* / *k*_*B*_ *T* (dimensionless when scaled)

###### Correspondence

- Embedding dimensions ↔ Molecular contact features
- Dot product ↔ Sum of contact energies
- Cosine similarity ↔ Normalized binding score

##### S4.7 Deterministic vs. Probabilistic Retrieval

###### S4.7.1 Top-k Retrieval (RAG Practice)

While softmax describes probabilistic retrieval, practical RAG uses deterministic top-*k*:

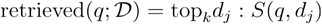

This returns the *k* documents with highest similarity.

###### Mathematical interpretation

Top-*k* is the *τ* → 0 limit of softmax retrieval (infinitely sharp selection).

###### S4.7.2 Threshold Activation (Immune)

Similarly, memory B cell activation involves thresholds:

- Cells with BCR affinity below threshold fail to activate
- Cells above threshold compete for T cell help

Both systems combine:

1. **Similarity-based ranking:** Score all candidates
2. **Threshold/top-k selection:** Activate best matches
3. **Competitive amplification:** Selected candidates produce output

##### S4.8 Resource Allocation

###### S4.8.1 Plasma Cell Niche Allocation

Bone marrow niches are limited. Which specificities get permanent slots?

###### Allocation depends on

- **Frequency:** Commonly encountered pathogens get more representation
- **Severity:** Dangerous pathogens get priority

###### Model

For pathogen *p* with frequency *f* _*p*_ and severity *s*_*p*_:

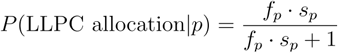

Complementary probability for memory B cell fate:

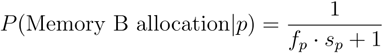

###### Interpretation

High-frequency, high-severity pathogens → LLPC; rare pathogens → memory B cells.

###### S4.8.2 Parametric vs. Retrieved Knowledge (RAG)

Similarly, RAG systems must decide what to encode where:

- **Frequently used knowledge:** Encode in parameters (faster access)
- **Rarely used knowledge:** Store in retrieval database (cheaper storage)
- **Critical knowledge:** May encode in both (redundancy)

###### Tradeoff

- Parametric: Fast, always available, limited capacity, expensive to update
- Retrieved: Unlimited capacity, requires retrieval step, cheap to update

##### S4.9 Why Dual Memory?

###### S4.9.1 The Fundamental Constraint

**You cannot have unlimited always-on memory.**

###### If everything were LLPCs / parametric

- Metabolic/compute cost prohibitive
- Bone marrow niches / model capacity overflow
- Updates require replacing existing memories

###### If everything were memory B cells / retrieved

- Response latency too slow for immediate threats
- Every response requires activation/retrieval overhead
- No baseline protection during retrieval delay

###### S4.9.2 Partitioning by Access Pattern

The dual architecture resolves this by partitioning:

| Access Pattern | Memory Type (Immune) | Memory Type (RAG) |
| --- | --- | --- |
| Frequently needed, time-critical | Plasma cells | Parametric |
| Rarely needed, can tolerate delay | Memory B cells | Retrieved |

##### S4.10 Summary

###### Structural correspondence

- Plasma cells ↔ Parametric memory (always-on, limited capacity)
- Memory B cells ↔ Document store (on-demand, large capacity)

###### Mathematical equivalence (retrieval/activation)

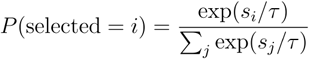

where *s*_*i*_ is BCR affinity or embedding similarity.

###### Strategic correspondence

- Dual architecture balances speed vs. capacity
- Similarity-based retrieval finds relevant memories
- Resource allocation prioritizes frequent/critical over rare

###### Derivation requires

- Immune memory biology (Ahmed & Gray, 1996)
- Plasma cell vs. memory B cell distinction (Slifka et al., 1998)
- BCR affinity-based activation (Paus et al., 2006)
- Boltzmann-form selection (same mathematics as Convergences 1–3)

##### S4.11 Verification Checklist

- [ ] LLPC properties accurately described
- [ ] Memory B cell properties accurately described
- [ ] Retrieval probability formula is correct softmax
- [ ] Top-k is limit of softmax as *τ* → 0
- [ ] Resource allocation model is reasonable
- [ ] Correspondence table entries are accurate
- [ ] Same Boltzmann form connects to previous convergences

## Supplementary Tables

**Supplementary Table S1.** Historical Development of Convergent Computational Strategies.

Historical Development of Convergent Computational Strategies
| Convergence | Component | AI Innovation | Seminal AI Reference | Immunological Concept | Seminal Immunology Reference |
| --- | --- | --- | --- | --- | --- |
| **1. Attention** | Core mechanism | Softmax attention over query-key | Vaswani et al. "Attention Is All You Need" (2017) | Boltzmann distribution governs binding | Boltzmann (1877); Amit et al. *Science* (1986) |
|  | Structural basis | Query-key-value decomposition | Vaswani et al. (2017) | Paratope-epitope contact structure | Amit et al. *Science* (1986) |
|  | Parallel channels | Multi-head attention | Vaswani et al. (2017) | Six CDR loops as independent recognition modules | Chothia & Lesk *J. Mol. Biol.* (1987) |
| **2. Contrastive** | Core mechanism | InfoNCE loss | Oord et al. *arXiv* (2018) | Clonal selection theory | Burnet (1959) |
|  | Selection probability | Softmax over similarities | Chen et al. *ICML* (2020) | Competitive selection dynamics | Kappler et al. *Cell* (1987) |
|  | Hard negatives | Hard negative mining | Robinson et al. *CVPR* (2021) | AIRE-mediated tissue-restricted antigen presentation | Anderson et al. *Science* (2002) |
| **3. Training** | Stage 1: Prior | Pre-training | Devlin et al. *NAACL* (2019) | Germline V(D)J recombination | Tonegawa *Nature* (1983) |
|  | Stage 2: Adaptation | Fine-tuning | Howard & Ruder *ACL* (2018) | Somatic hypermutation | Weigert et al. *Nature* (1970) |
|  | Stage 3: Calibration | RLHF | Ouyang et al. *NeurIPS* (2022) | T follicular helper selection | Breitfeld et al. *J. Exp. Med.* (2000) |
| **4. Memory** | Dual architecture | RAG | Lewis et al. *NeurIPS* (2020) | Plasma cells + Memory B cells | Ahmed & Gray *Science* (1996) |
|  | Always-on memory | Parametric weights | — | Long-lived plasma cells | Slifka et al. *Immunity* (1998) |
|  | On-demand memory | Document retrieval | Karpukhin et al. *EMNLP* (2020) | Memory B cell reactivation | Kurosaki et al. *Nat. Rev. Immunol.* (2015) |

**Supplementary Table S2:**
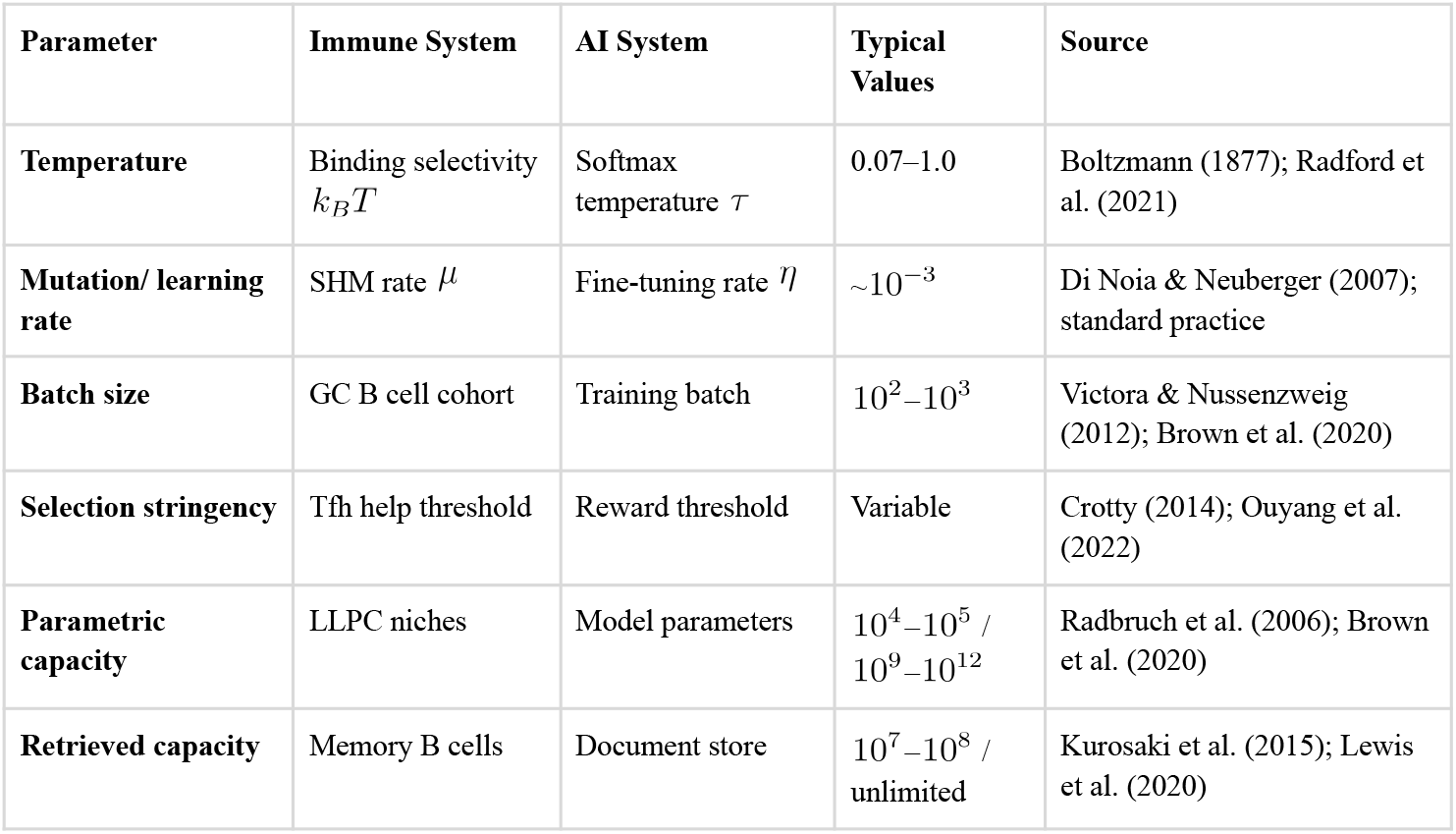
Parameter Correspondences.

**Supplementary Table S3:**
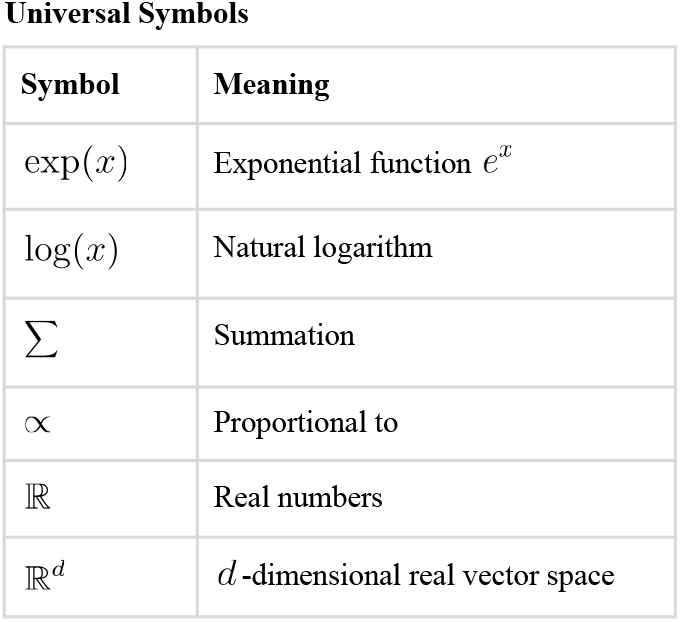
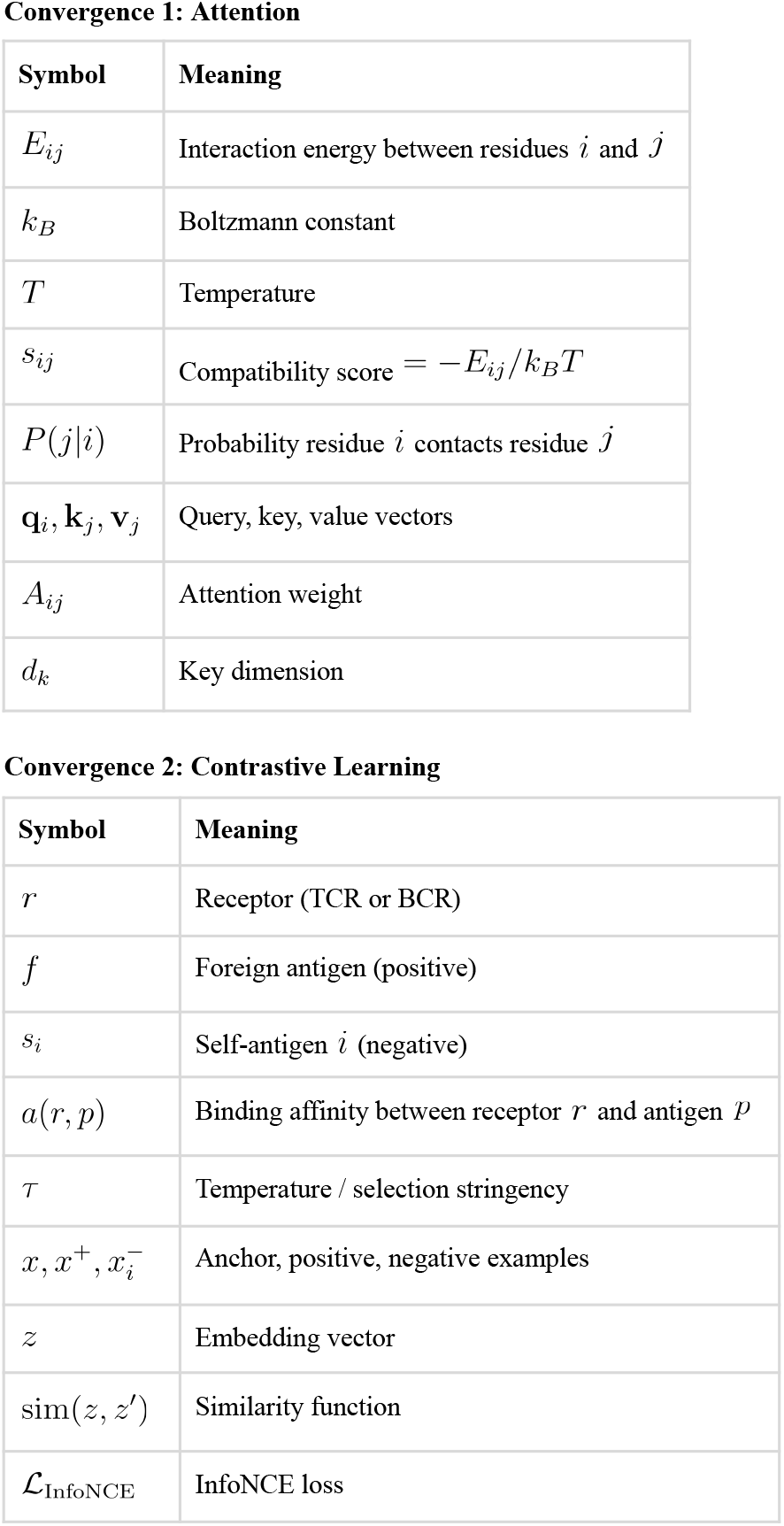
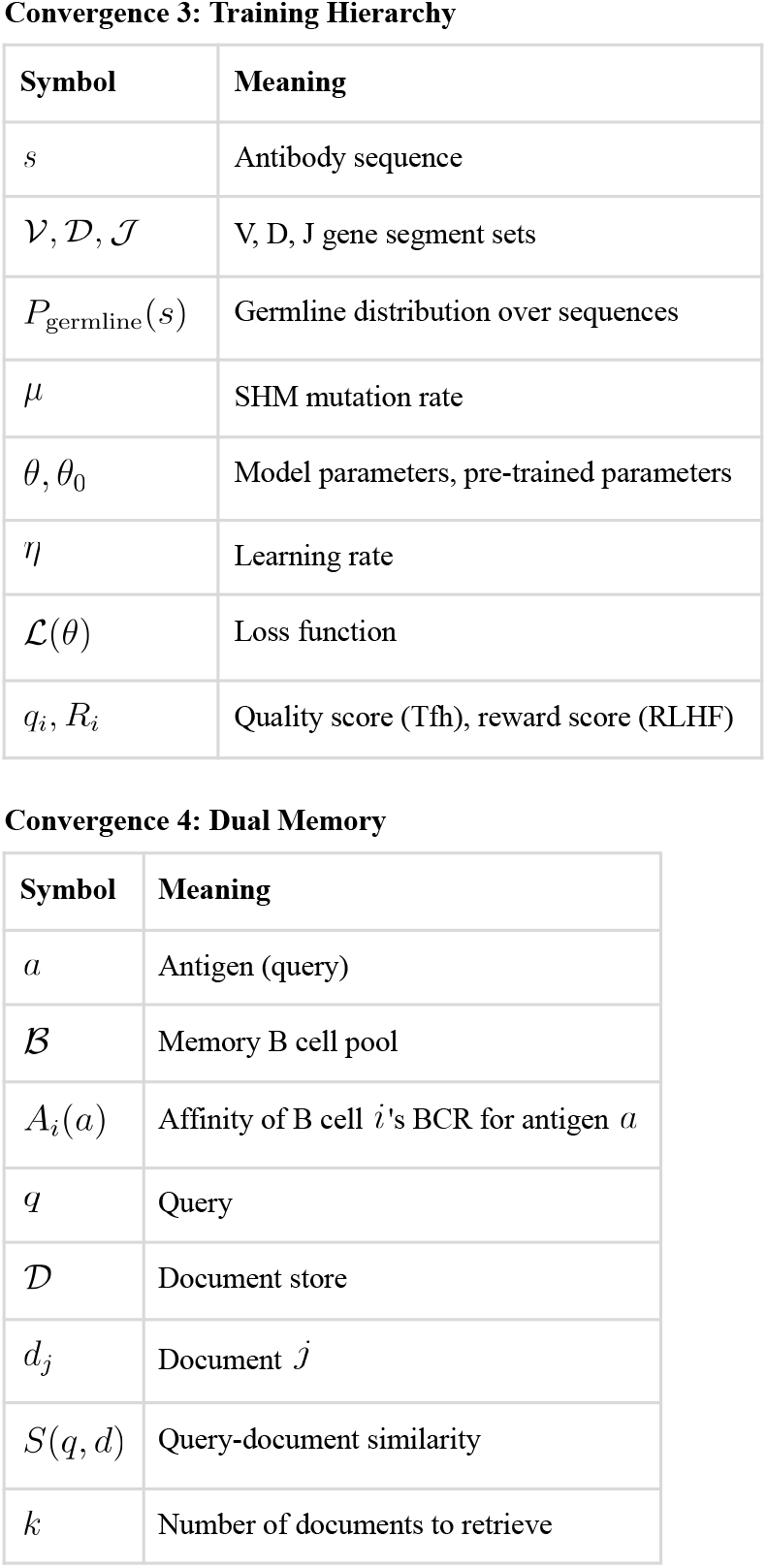
Notation Glossary Universal Symbols.

| Symbol | Meaning |
| --- | --- |
| $\exp(x)$ | Exponential function $e^x$ |
| $\log(x)$ | Natural logarithm |
| $\sum$ | Summation |
| ∝ | Proportional to |
| $\mathbb{R}$ | Real numbers |
| $\mathbb{R}^d$ | $d$-dimensional real vector space |

**Convergence 1: Attention**
| Symbol | Meaning |
| --- | --- |
| $E_{ij}$ | Interaction energy between residues $i$ and $j$ |
| $k_B$ | Boltzmann constant |
| $T$ | Temperature |
| $s_{ij}$ | Compatibility score $= -E_{ij}/k_B T$ |
| $P(j\vert i)$ | Probability residue $i$ contacts residue $j$ |
| $\mathbf{q}_i, \mathbf{k}_j, \mathbf{v}_j$ | Query, key, value vectors |
| $A_{ij}$ | Attention weight |
| $d_k$ | Key dimension |

**Convergence 2: Contrastive Learning**
| Symbol | Meaning |
| --- | --- |
| $r$ | Receptor (TCR or BCR) |
| $f$ | Foreign antigen (positive) |
| $s_i$ | Self-antigen $i$ (negative) |
| $a(r, p)$ | Binding affinity between receptor $r$ and antigen $p$ |
| $\tau$ | Temperature / selection stringency |
| $x, x^+, x_i^-$ | Anchor, positive, negative examples |
| $z$ | Embedding vector |
| $\text{sim}(z, z')$ | Similarity function |
| $\mathcal{L}_{\text{InfoNCE}}$ | InfoNCE loss |

**Convergence 3: Training Hierarchy**
| Symbol | Meaning |
| --- | --- |
| $s$ | Antibody sequence |
| $\mathcal{V}, \mathcal{D}, \mathcal{J}$ | V, D, J gene segment sets |
| $P_{\text{germline}}(s)$ | Germline distribution over sequences |
| $\mu$ | SHM mutation rate |
| $\theta, \theta_0$ | Model parameters, pre-trained parameters |
| $\eta$ | Learning rate |
| $\mathcal{L}(\theta)$ | Loss function |
| $q_i, R_i$ | Quality score (Tfh), reward score (RLHF) |

**Convergence 4: Dual Memory**
| Symbol | Meaning |
| --- | --- |
| $a$ | Antigen (query) |
| $\mathcal{B}$ | Memory B cell pool |
| $A_i(a)$ | Affinity of B cell $i$'s BCR for antigen $a$ |
| $q$ | Query |
| $\mathcal{D}$ | Document store |
| $d_j$ | Document $j$ |
| $S(q, d)$ | Query-document similarity |
| $k$ | Number of documents to retrieve |

